# Mosaic representations of odors in the input and output layers of the mouse olfactory bulb

**DOI:** 10.1101/259945

**Authors:** Honggoo Chae, Daniel Kepple, Walter G. Bast, Venkatesh N. Murthy, Alexei Koulakov, Dinu F. Albeanu

**Affiliations:** Cold Spring Harbor Laboratory, Cold Spring Harbor, NY; Watson School for Biological Sciences, Cold Spring Harbor, NY; Department of Molecular & Cellular Biology and Center for Brain Science, Harvard University, Cambridge, MA

**Author notes:** equal contribution (Honggoo Chae, Daniel Kepple).

**Keywords:** glomeruli, mitral and tufted cells, intrinsic signals, two photon calcium imaging, distributed representation, topography, odor space, physical-chemical odor properties, LASSO

## Abstract

The elementary stimulus features encoded by the olfactory system remain poorly understood. We examined the relationship between 1,666 physical-chemical descriptors of odors and the activity of olfactory bulb inputs as well as outputs in awake mice. Glomerular and M/T cell responses were sparse and locally heterogeneous, with only a coarse dependence of glomerular positions on physical-chemical properties. Odor features represented by ensembles of M/T cells were overlapping, but distinct from those represented in glomeruli, consistent with extensive interplay between feedforward and feedback inputs to the bulb. This reformatting was well-described as a rotation in odor space. The descriptors accounted for a small fraction in response variance, and the similarity of odors in physical-chemical space was a poor predictor of similarity in neuronal representations. Our results suggest that commonly used physical-chemical properties are not systematically represented in bulbar activity and encourage further search for better descriptors of odor space.

## Introduction

Insight into the computations performed by a sensory system often arises from knowing which features of inputs from the environment are encoded by the underlying neuronal circuits - for example, wavelength of light or frequency of sound^1,2^. Unlike for other sensory modalities (i.e. vision, audition), it is not understood what properties of odors are important in olfaction, how these properties are processed by olfactory neural circuits, and how odorant receptor genes have evolved to optimize such an encoding. The relationship between odor structure (chemical space), the underlying spatial and temporal patterns of activity in the brain (neuronal space) and the perceived odor quality (perceptual space) has been elusive^3-12^. As a result, a basic goal remains unachieved: to robustly predict neuronal activity and perceptual attributes, starting from the physical features of odor molecules.

Even the earliest step in olfaction, the interaction between a particular odorant receptor (OR) and ligands (odorants) has defied simple descriptions. Additional complexity arises from the numerous distinct OR types present in the olfactory system of many vertebrates, leading potentially to a large number of dimensions of the odor space that could be sampled by the brain. Recent studies, enabled by advances in computing power, have used dimensionality reduction strategies to suggest that a relatively small number of odor physical-chemical descriptors (~30 out of ~1,600 to ~5,000 in the eDragon database^13^) capture odor similarity in all three spaces - chemical, perceptual and neuronal^7,14-21^. In addition, it has been proposed that most variance in human perceptual space can be explained by considering low dimensional manifolds (~2-20) along orthogonal axes that are correlated to behaviorally relevant features such as stimulus pleasantness, toxicity and/or hydrophobicity ^4,7,9,10,16,17,22,23^.

The spatial layout and wiring patterns of neural circuits can offer important clues about the underlying computations - for example, a retinotopic organization along with nearest neighbor interactions allowed the inference of local contrast enhancement in the retina^24,25^. In the vertebrate olfactory system, there is a predictable and precise layout of OR identity in the glomerular layer of the OB^26,27^. However, if, or how this receptor-based layout translates to a functional map remains unclear^18,28-35^.

Odor information arriving at glomeruli from the olfactory sensory epithelium is modified within the olfactory bulb (OB) by local and top-down interactions. Two classes of output neurons, mitral and tufted cells (M/T), convey information to several olfactory cortical areas. These two output channels differ in their inputs, morphology, intrinsic excitability, local connectivity, activity patterns, downstream targets and top-down feedback^36-40^.

Inspired by recent progress in relatively unbiased characterization of odors through the use of a large number of physical-chemical properties, we set out to investigate whether these features are represented as systematic and recognizable spatial-temporal neuronal activity patterns in the input and/or output layers of the bulb. We studied how the activity patterns of glomeruli and M/T cells in awake, head-fixed mice relate to a set of commonly used 1,666 physical-chemical odor properties. Using intrinsic optical imaging, multiphoton microscopy, dimensionality reduction and regression techniques, we asked four general questions. First, how well are these physical-chemical properties of odors captured by the glomerular and M/T cells odor responses? Second, does similarity of odor molecules in the physical-chemical properties space correlate with the neuronal population responses? Third, does the physical location of individual glomeruli or M/T cells depend on their physical-chemical receptive fields? Finally, what is the relationship between neuronal odor representations in the input and output layers of the bulb?

## Results

We used imaging of intrinsic signals from the dorsal surface of the olfactory bulb to probe the responses of glomeruli to an array of chemically diverse odors (Supplementary Figure 1, Supplementary Table 1, Methods). Previous reports indicate that glomerular intrinsic signals approximate well the activity of presynaptic OSN terminals in the olfactory bulb^29,41,42^. In separate experiments, we employed multiphoton microscopy to monitor the responses of mitral and tufted cells tilling the dorsal aspect of the bulb to the same stimuli (Supplementary Figures 1,2,3) across a range of concentration (Supplementary Note 1, Methods). We used mice that express the genetically encoded calcium indicator GCaMP3 in mitral and tufted cells (by crossing TBET-Cre with Ai38, floxed GCaMP3 animals^43^). For both glomerular (n=6 mice, 12 bulb hemispheres/fields of view) and M/T cells (n=15 mice, 19 fields of view) imaging sessions, we identified regions of interests (ROIs) that were responsive to the odors in our panel (1,057 glomeruli and 1,537 M/T somata, Supplementary Note 1, Supplementary Figures 1-3, Methods).

### Odor responses of glomeruli and M/T cells are poorly described by eDragon physical-chemical properties

Previous studies suggested that the olfactory system is tuned to extract specific physical-chemical features of odors^3,9,10,15,16^. Here, we investigated whether a wide array of physical-chemical molecular parameters are good predictors of glomerular and M/T cell responses. We used the eDragon database^13^ to assign values for 1,666 physical-chemical properties to each odor in the panel. The responsiveness of each ROI (glomerulus or M/T cell), or the property response (PR), to a particular property was characterized as a Pearson correlation between the odor responses of the ROI and the values taken by the property (property strength vector, PSV) across the odors in the panel (Figure 1a-c, Methods). 49 different monomolecular odors were used in both the glomerular and M/T cell imaging experiments. The array of such correlations across the eDragon molecular properties is defined as the property response spectrum (PRS) of that glomerulus or M/T cell (Figure 1d,g). This definition is analogous to calculating a neuronal receptive field in the visual or auditory systems, where the response strength of a given cell is correlated with certain stimulus features. The PRS represents the physical-chemical properties that a given ROI responds to, thus playing the role of a molecular receptive field.

**Figure 1.**
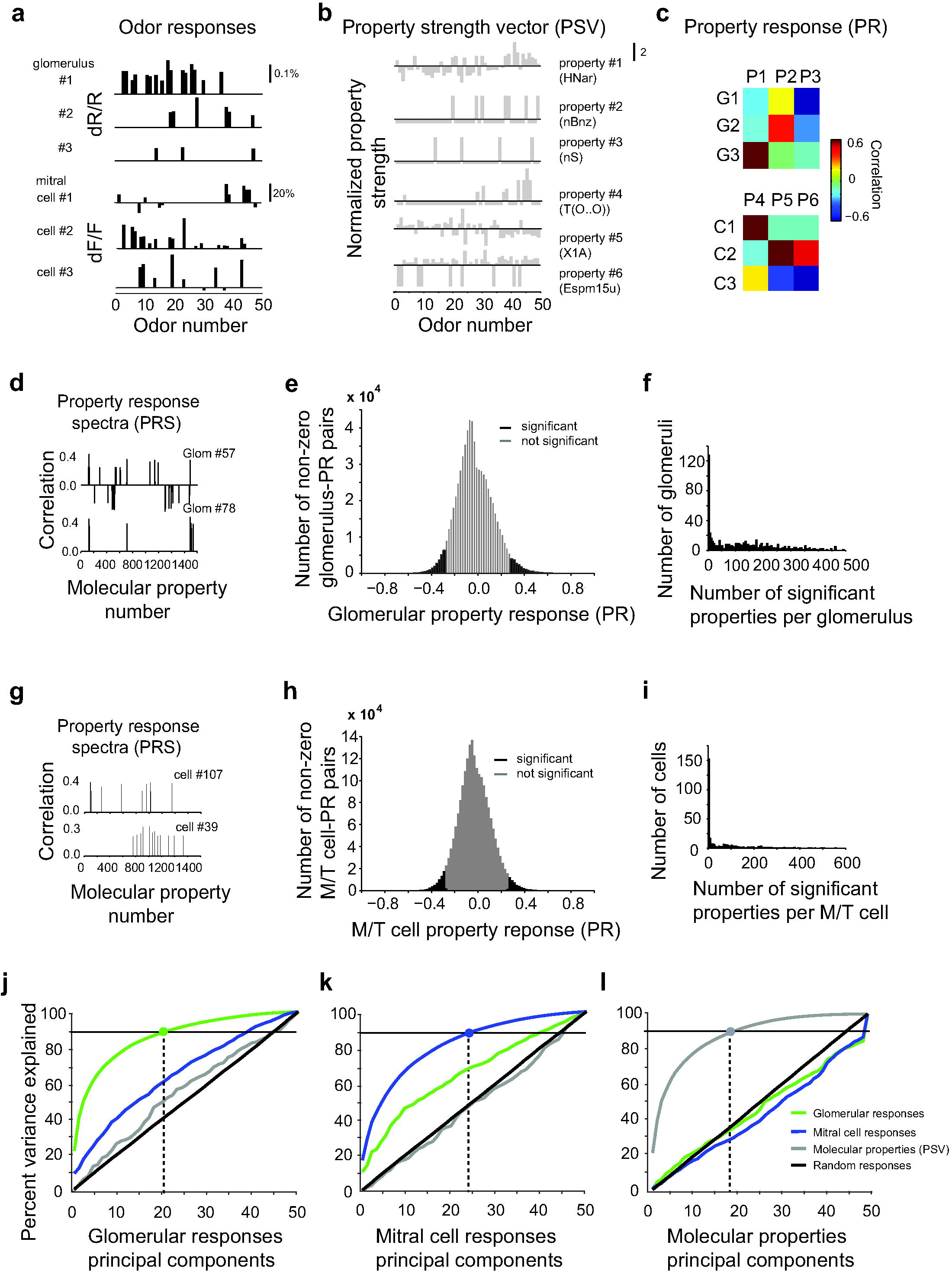
Tuning of glomeruli and mitral/tufted cells responses to eDragon physical-chemical properties. **a**. Odor responses of three example glomeruli and mitral cells. The responses (dR/R, dF/F) are shown for a panel of 49 monomolecular odors. Non-significant responses were set to zero (Methods). **b**. Six example molecular property strength vectors (PSV) across same odors as in (**a**); HNar (Narumi harmonic topological index), nBZ (number of benzene rings), nS (number of sulfur atoms), T(O̤O) (sum of topological distances between oxygen atoms), X1A (average connectivity index of order 1), Espm15u (spectral moment of order 15 from edge adjacency matrix). **c**. Property responses (PR) given by the Pearson correlation coefficients between the odor responses (**a**) of the three example glomeruli (*top*) and mitral cells (*bottom*), and the 6 example molecular property strength vectors (**b**) calculated over 49 odors in the panel. **d,g**. Example correlations (property response spectra, PRS) between odor responses and molecular properties for two example glomeruli (**d**) and mitral cells (**g**). **e, h**. Distribution of the glomerular response-molecular property (**e**) and M/T cell response-molecular property (**h**) pairwise correlations. Significant correlations are shown in black and nonsignificant ones in grey (Methods). **f, i.** Histogram of the number of molecular properties that individual glomeruli (**f**) and M/T cells (**i**) respond to, above the significance threshold, with an average of 62 per glomerulus and 67 per M/T cell. **j, k, l**. Results of principal component analysis, PCA for glomerular (**j**), M cells responses (**k**) and molecular properties (**l**). Percent of variance explained is shown as a function of the number of included principal components, PCs. (**j**) Percent variance explained of glomerular (green), mitral cells (blue) odor responses, molecular property strength vectors (grey) and random data controls (black) shown as function of the number of included glomerular responses principal components. (**k,l**) Percent variance explained of glomerular (green), mitral cells (blue) odor responses, molecular property strength vectors (grey) and random data controls (black) shown as function of the number of included mitral cell responses principal components (**k**) and of molecular properties principal components (**l**).

For both glomeruli and M/T cells, the number of properties that single units responded to significantly (Methods) varied from ROI to ROI, with an average of 62 per glomerulus and 67 properties per M/T cell (Figure 1f, i). In general, individual responsive glomeruli and M/T cells were only poorly tuned to the physical-chemical properties (average correlation of −0.02±0.16 SD for glomeruli and −0.02±0.15 SD for M/T cells, Supplementary Figure 4a,c; ~26% of glomeruli and ~33% of M/T cells showed no significant correlations to any of these properties), and the vast majority of glomerular and cell odor response-property correlation pairs was not significant (Figure 1d-i, Methods). Within the subset of significant glomerular and cell odor response-property pairs (~6%), correlations spanned both negative and positive values (absolute average: 0.35±0.07 SD for glomeruli and 0.36±0.08 SD for M/T cells, Supplementary Figure 4e). Across different experiments, the tuning widths (number of significant properties per ROI) were similar (Supplementary Figure 4b,d), indicating similarities in the overall tuning to different molecular properties on the dorsal surface of the bulb across animals.

We further used principal component analysis (PCA) to quantify the dimensionality of three different datasets that relate the physical and neuronal odor representations (Figure 1j,k,l). Approximately 17 dimensions were sufficient to account for 90% of variance in the values taken by these molecular properties across the 49 odors used (Figure 1l). Thus, many of these properties are redundant^15,16^, and a 17D flat surface contains substantial amount of information (90%) on the molecular properties. The dimensionality of the physical-chemical descriptors depends on the odors included in the panel, and, in principle, can be independent of the responses of olfactory neurons. In comparison, the responses of glomeruli to odors in our panel could be described within a 21D principal components (PC) space, and those of the M/T cells in a 24D space at the same level of variance explained (90%, Figure 1j,k).

Are these molecular properties efficiently represented by the neural responses? To answer this question, we used a computational technique that we call principal component exchange (PCX) method (Methods). We projected the molecular properties (property strength vectors) into the space of glomerular responses principal components. We then measured the variance of molecular properties data captured by the glomerular responses PC space of increasing dimensionality (Figure 1j). If the molecular and neuronal responses PC spaces were identical, then glomerular responses PCs would capture the same percent variance in both glomerular responses and the molecular properties datasets. Instead, we find that glomerular PCs explain almost the same fraction of variance in the molecular properties dataset as they do in randomly generated data (Methods, grey and black lines, Figure 1j). Mitral cell responses PCs also captured a similar fraction of variance in the molecular properties as in the random dataset (grey and black lines, Figure 1k). When projected to the PC space of molecular properties, glomerular and mitral (same for tufted, data not shown) cell responses, were similar to random data controls as well in terms of amount of variance explained (Figure 1l). Projecting mitral cell responses in the glomerular PCs space, or glomerular responses in the mitral cells PC space captured less variance compared to the reference PC spaces (glomerular and mitral), but substantially more than random data (discussed later, together with a comparison between mitral and tufted cell responses, Figure 5). Taking together these results, indicate that the molecular properties, as defined by the eDragon descriptors, are underrepresented in the glomerular and M/T cell odor responses.

Overall, our data indicates that neural responses of both glomeruli and olfactory bulb outputs are poorly tuned to the physical-chemical properties analyzed, and instead reflect features of the odors that are not well captured by these molecular properties commonly used in computational chemistry, and by previous studies in olfaction.

### Odor similarity in physical odor space is a poor predictor of neuronal representations

Does the similarity between pairs of odors, calculated using the set of 1,666 physical-chemical properties reflect the similarity in neuronal representations of the same odors in either the input or output layers of the OB? To describe similarity in odor physical space, for each odor pair in the panel, we calculated the Euclidean distance between the normalized molecular properties strength vectors (PSV) associated with each odor in the panel (Figure 1b) as proposed by previous studies^15,16,18,44^. To represent odor similarity in the neuronal representations, we used two metrics: 1) the Euclidean distance between the ROIs responses to the same pair of odors, and 2) the Pearson correlation in neuronal responses (pooling data across fields of view, Methods). For both glomeruli and M/T cells, the pairwise odor similarity in the space defined by the physical-chemical properties had only poor and variable correlation with the similarity in neuronal representations (Figure 2a, b). Therefore, neuronal responses in the bulb may not faithfully represent the multidimensional vectors of odor properties.

**Figure 2.**
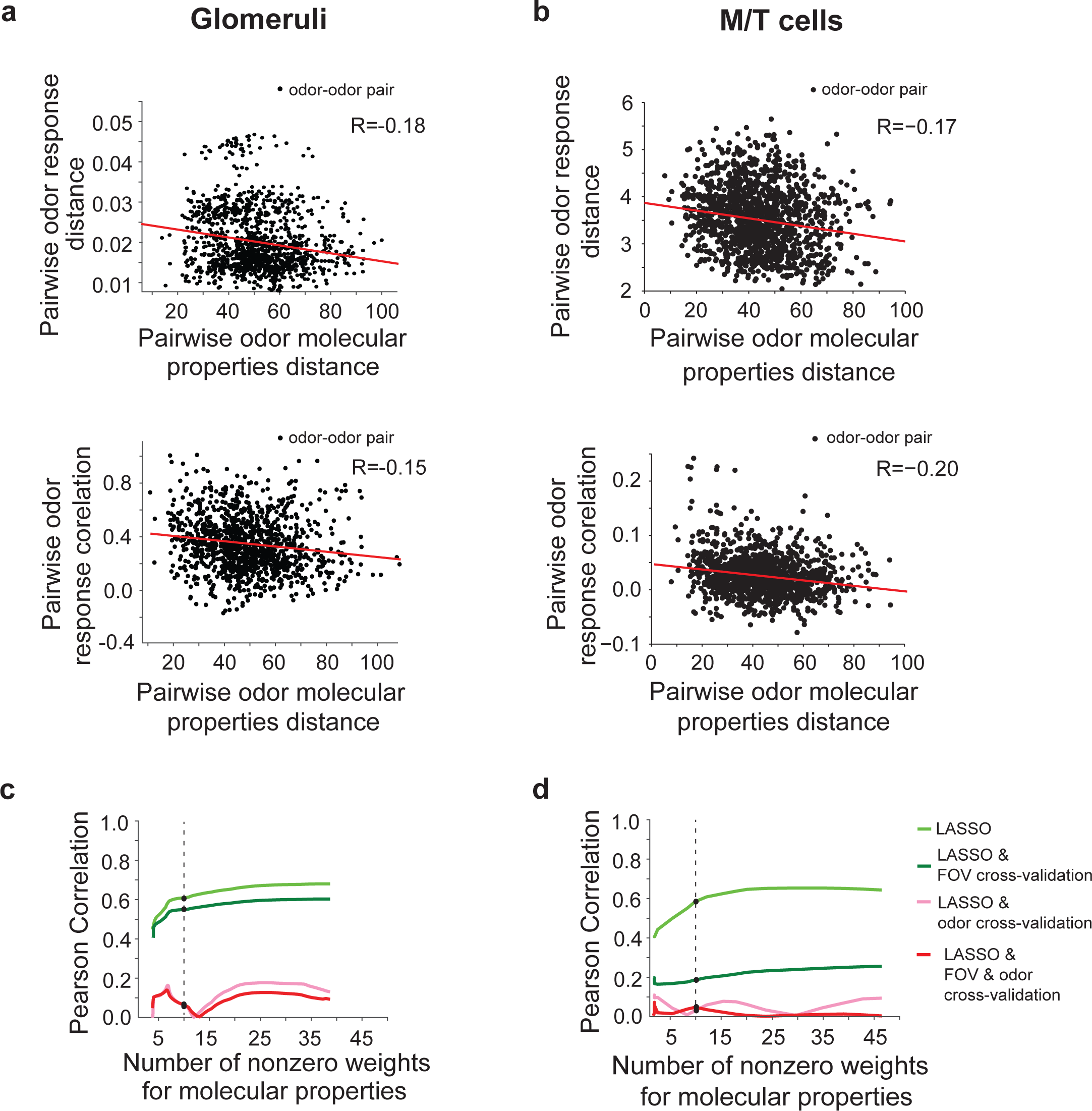
Pairwise odor similarity comparison across physical-chemical and neuronal response odor representations. **a**. Pairwise odor Euclidean distance across eDragon physical-chemical properties versus distance between glomerular responses, expressed as the Euclidean distance (R=-0.18, *Top*) and Pearson correlation (R=-0.15, *Bottom*). **b**. Pairwise odor Euclidean distance across eDragon physical-chemical properties versus distance between M/T cell responses, expressed as the Euclidean distance (R=-0.17, *Top*) and Pearson correlation (R=-0.20, *Bottom*). Note that average odor response Euclidean distances for glomeruli and M/T cells representations are expected to differ since they are determined by the absolute strength of intrinsic and fluorescence signals. **c**. Least absolute shrinkage and selection operator (LASSO) regression based on a subset of physical-chemical properties selected from the 1,666 set, describing the relationship between odor pairwise similarity across properties versus glomerular responses; (*light green*) all imaged hemibulbs were used as a training set for optimizing the regression; (*darkgreen*) half of the fields of view were used as a training set and the remaining half for cross-validation (FOV cross-validation); (*pink*) all imaged hemibulbs were used for training, while one pair of odors was iteratively left out during training and added back subsequently for cross-validation (jackknife, odor cross-validation); (*red*) half of the fields of view were used for training and the remaining half for cross-validation; in addition, one pair of odors was iteratively left out during training and subsequently added back for cross-validation (jackknife, FOV and odor cross-validation). **d**. Least absolute shrinkage and selection operator (LASSO) regression based on a subset of physical-chemical properties selected from the 1,666 set, describing the relationship between odor pairwise similarity across properties versus M/T cell responses; (*light green*) all FOVs of imaged M/T cells were used as a training set for optimizing the regression (R=0.68); (*darkgreen*) half of the fields of view were used as a training set and the remaining half for cross-validation (FOV cross-validation, R=0.2); (*pink*) all imaged FOVs of mitral and tufted cells were used for training, while one pair of odors was iteratively left out during training and added back subsequently for cross-validation (jackknife, odor cross-validation, R=-0.06); (*red*) half of the fields of view were used for training and the remaining half for cross-validation; in addition, one pair of odors was iteratively left out during training and subsequently added back for cross-validation (jackknife, FOV and odor cross-validation, R=-0.08); dotted line corresponds to an instantiation of the LASSO regression, when only 10 physical-chemical properties were allowed took non-zero weights;

Perhaps, only a small subset of the molecular properties has a robust relation to the neural responses. To determine whether a particular subset of molecular properties is more informative of neuronal responses, for each odor pair, we built a sparse regression between the squared neuronal pairwise odor response distances and the squared differences between individual physical-chemical property values. We used a non-negative Least Absolute Shrinkage and Selection Operator (LASSO) algorithm that selects a sparse subset of non-zero properties from the full set to better explain the differences in neuronal responses^45^ (Methods). The properties were selected such as to yield the best fit of pairwise odor distances in neuronal responses and physical-chemical space, by weighting the odor similarity calculated in the physical-chemical space. If all molecular property weights had values of 1, one would arrive at the correlation values shown in Figure 2a and b. To find a sparse and robust solution, the LASSO algorithm assigns zero value to the weights of most molecular properties. By varying a penalty parameter (*λ*) of the algorithm (Methods, Supplementary Figure 4f), we changed the number and relative contribution of the molecular properties included in computing the pairwise odor distances. Using this approach, we could identify small subsets of properties which are well reflected in the neuronal responses. For example, using 10 molecular properties selected via LASSO, the correlation between the pairwise odor distances in physical-chemical and neuronal responses increased substantially (~0.60); we ran LASSO independently for the glomerular and M/T cell data sets, Methods). Including more properties into the analysis mildly improved the correlation (40 properties brought the correlation score up to ~0.65, Figure 2c,d).

We further determined how these regression analyses and correlations generalize across different FOVs (FOV cross-validation), and odor pairs (odor cross-validation). In individual iterations of the FOV cross-validation analysis, we randomly removed half of the FOVs and performed the sparse linear regression of properties on the remaining FOVs (training set). The best fit obtained was then used to predict the neuronal space odor similarity for the removed FOVs. In individual iterations of the odor cross-validation analysis, we removed one pair of odors from the panel (jackknife, leave one out), performed the sparse linear regression on the remaining odors, and further used it to predict the neuronal space similarity for the removed odor pair. This procedure was repeated independently for each odor pair. Thus, every prediction of an odor pair similarity was obtained based only on odor similarities calculated for all other odor pairs. Crossvalidation across fields of view (FOV) decreased only slightly the observed glomerular responses-to-physical space correlations, consistent with reproducible glomerular odor maps across individuals^29^. The same procedure however resulted in substantially lower correlations (~0.2) for the M/T cell responses, which may reflect differences in sampling M/T cells along the dorsal aspect of the OB across different fields of view. In contrast, the odor cross-validation procedure, or the combination of these two procedures, drastically diminished the correlations to the physical-chemical representations (at most ~0.10 for glomeruli and ~0.05 for M/T cells, Figure 2c,d, Methods). In addition, employing a greedy algorithm (Methods) with odor cross-validation for the M/T cells data led to qualitatively similar results (Supplementary Figure 4g).

Thus, while correlations between similarity in physical-chemical and neuronal response spaces could be identified using subsets of molecular properties, they held little predictive power when new odor pairs were tested, suggesting that such correlations emerge due to over fitting.

### Relating the tuning of glomeruli and M/T cells to molecular properties to their placement on the bulb

Previous experiments in the glomerular layer of the rodent olfactory bulb have provided contrasting views of spatial organization. Some authors have suggested that different classes of chemicals are represented in a spatially segregated manner, perhaps even in an ordered topographic manner^34,35,46-51^. Others have noted a great deal of local disorder, with no evidence of a smoothly-varying representation^18,29,52^. To date, these spatial relations have not been assessed using physical-chemical properties of odors, such as those provided by eDragon. Although our analysis above has shown that glomeruli and M/T cells sample a small subspace of the properties (Figure 1), some of these parameters are represented in the neuronal responses. Therefore, we investigated whether tuning to the molecular properties is spatially laid out in a systematic fashion at the level of glomeruli and M/T cells.

For each property, we characterized the relationship between the tuning of glomerular and M/T cells responses, and the spatial location of glomeruli/cells along the anterior-posterior (AP) and medial-lateral (ML) axes of the bulb. The tuning of individual ROIs (glomeruli or M/T cells) was described by their property responses (PR). For the ensemble of ROIs monitored in each FOV, we computed the correlation between each ROI’s location, determined experimentally, and its response sensitivity to each property. A significant correlation between the response sensitivity to a given property (PR) and an ROI location in a field of view would suggest that the glomeruli/M/T cells in that FOV are spatially organized by their sensitivity to that property. For each FOV (2D plane), every molecular property is described by the strength of its correlation along the AP and ML axes. The significance of these correlations has to be analyzed carefully, as we test for a large number of molecular properties in each FOV. We therefore implemented a false discovery rate (FDR) correction^53^ to control the rate of false positives (Figure. 3a-f, 4a,b, Methods).

**Figure 3.**
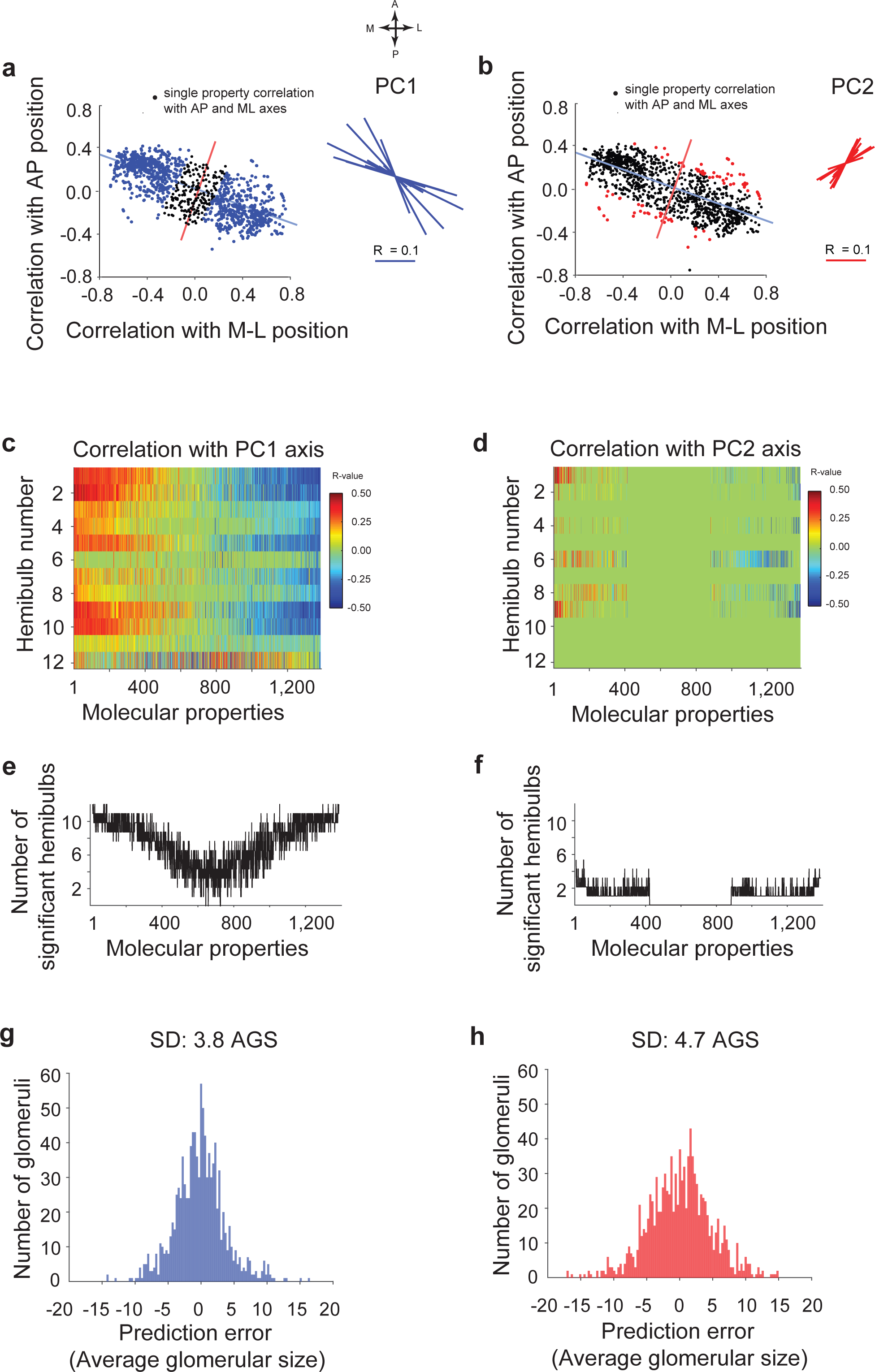
Coarse spatial correlations between glomerular positions and molecular property responses. **a, b**. Pearson correlations between individual property responses (PR) and the placement of glomeruli along the anterior-posterior (AP) and medial-lateral (ML) axes of the bulb. Each dot represents the correlation value of an individual molecular property. Performing PCA on the cloud of molecular property correlations computes the orientation of the first principal component (PC1, blue) with respect to the AP and ML anatomical axes. Properties shown by blue and black dots correspond to significant and non-significant correlations with the PC1 axis (FDR q<0.1). Red dots show properties significantly correlated with PC2 axis (orthogonal on PC1, FDR q<0.1). Orientations of the PC1 and PC2 axes in 12 hemibulbs (bulb hemispheres) are shown by blue and red lines. **c, d**. Correlations between individual molecular properties and glomerular positions along the PC1 and PC2 axes for each hemibulb. In each panel, properties are re-sorted with respect to the strengths of correlation. For the panel of odors used, several properties (~200) did not take nonzero values, and were not included in the analysis. **e, f.** Number of hemibulbs in which a given molecular property is significantly correlated with the glomerular position along PC1 (**e**) and PC2 (**f**). A set of properties is consistently correlated with the PC1 axis (at most 11 out of 12 hemibulbs). Correlations along PC2 are less consistent across samples (at most 2 out of 12 hemibulbs). **g, h**. Histogram of normalized displacement error vectors between the location of observed and predicted glomerular locations along PC1 and PC2 axes. The predictor was obtained using a LASSO algorithm (jackknife cross-validation) to build a sparse linear regression based on 20 molecular properties (Methods). Prediction error is shown along PC1 (*Left*, SD=3.8 average glomerular size, AGS) and PC2 axes (*Right*, SD=4.7 AGS).

**Figure 4.**
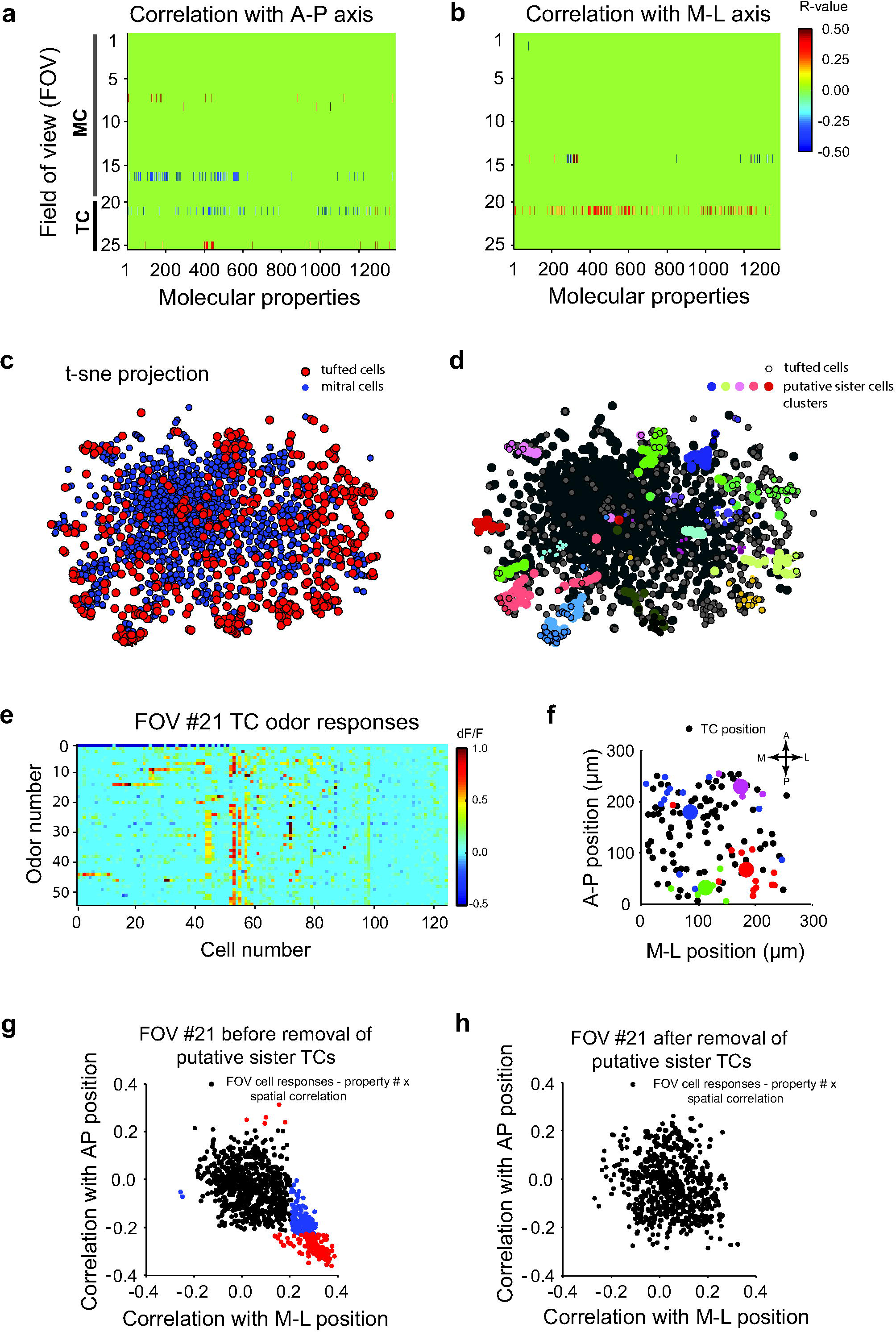
Lack of correlation between mitral/tufted cell placement and physical-chemical odor properties. **a, b**. Pearson correlation coefficients between M/T cell locations with respect to the anterior-posterior (AP, *Left*) and medial lateral (ML, *Right*) axes within fields of view, FOVs (rows) and their response sensitivity to individual properties across the physical-chemical properties analyzed. Only significant correlations are shown (FDR, q<0.1). FOVs were sorted for mitral and respectively tufted cells for both nominal dilutions (1:100 and 1:3,000). **c**. t-sne projection of responses of all sampled mitral (blue) and tufted (red) cells. Each dot represents an individual M/T cell from one of the 19 FOVs imaged. **d**. Same t-sne projection as in (**c**) with different putative co-glomerular clusters shown by different colors. Tufted cell responses are represented by black circles. **e**. Odor responses (dF/F) of tufted cells from the example FOV (#21) sorted by size of functional clusters (Methods). Horizontal bars (Top) mark clusters of putative co-glomerular sister cells. **f**. Positions of tufted cells in the example FOV. Color dots mark clusters of putative sister cells. Larger colored dots correspond to the average location of putative sister cells within a given functionally defined cluster. **g, h**. Example field of view illustrating the correlation of physical-chemical properties with placement of cells along the anterior-posterior (AP) and medial-lateral (ML) axes of the bulb before (**g**) and after (**h**) removal of putative co-glomerular sister cells via odor tuning based clustering. Each point represents an individual molecular property. Properties significantly correlated (FDR q<0.1) with either AP or ML axes are displayed as blue and red dots respectively; black dots represent non-significant correlations.

When evaluating the layout of molecular properties in space, it is important to realize that the AP and ML axes are not special. Other directions on the bulb surface could be well aligned with independent combinations of the odor physical-chemical properties. To discover whether any such particular axes exist, we first performed PCA on the glomerular data, thus rotating the coordinates such that the combinations of molecular properties correlated with each of the two new reference axes (PC1 and PC2) are independent from each other (Figure 3 a, Methods). Across different animals, the first principal axis (PC1, blue, Figure 3) was consistently rotated approximately 40 degrees (Avg=38.7±16.3 SD) with respect to the A-P direction. Several molecular properties (20) appeared strongly correlated (positively and negatively) with PC1 (Figure 3c,e, Supplementary Figure 5, Supplementary Table 2), and could be identified robustly across animals (11 out of 12 hemibulbs). The properties correlated with the PC2 axis were much less consistent across samples (2 out of 12 hemibulbs, Figure 3d,f). The orientation of the principal axis (PC1) we identify here may reflect specific interactions between the axon terminals of OSNs and gradients of axon guidance molecular cues during the formation of the glomerular map (Discussion).

### The responses of glomeruli are predictive of their location on the bulb surface at a coarse spatial scale

Could the positions of individual glomeruli on the bulb be inferred using the properties they are responsive to? We tested this hypothesis by building a sparse linear regression for individual glomerular positions based on their tuning (PRS) to the molecular properties. Regression was obtained as above, using a LASSO algorithm that selects a small subset of active properties (20) from the full set (Methods). Different sets of properties were selected such as to yield the best match of glomerular positions for each FOV and reference axes (PC1 vs. PC2). We verified that changing the number of active properties does not affect our results substantially and tested the quality of our predictions using a jackknife cross-validation. To quantify the quality of prediction, we evaluated the prediction error for each glomerulus for PC1 and PC2 normalized by the average glomerulus spacing (AGS)^29^. We find that glomerular positions are defined more precisely along the PC1 versus PC2 axis (SD = 3.8 versus 4.7 AGS, Figure 3g,h). We also performed the same analysis using the odor response spectra of individual glomeruli instead of their sensitivity to the molecular properties (PR), and reached quantitatively similar results (data not shown).

Importantly, shuffled controls (obtained by randomizing the identity of molecular properties) produced statistically indistinguishable outcomes in the prediction of glomerular positions by using shuffled property responses along newly computed PC1 and PC2 axes (data not shown). This is somewhat expected, given our strict selection procedure that relies on LASSO to identify the best small set of properties (20) out of a much larger pool such as to minimize the error between the observed and predicted positions.

Overall, consistent with earlier work^29,50^, these results indicate that responses of glomeruli to odors and, in particular, to a set of molecular properties are moderately predictive of their coarse placement on the olfactory bulb surface.

### Tuning of M/T cells to molecular properties is not correlated with their spatial location

To investigate the relationship between location of OB output neurons and their tuning to molecular properties, we carried out the same analysis on the responses of M/T cells to the 49 monomolecular odors. In general, M/T cell responses in a FOV were locally heterogeneous, responding to chemically diverse odors (Supplementary Note 2). The resulting correlations are represented as a raster plot for all FOVs in Figure 4a,b. Most sampled FOVs showed very few or no significant correlations at all. Strikingly, in one FOV (#21, tufted cells), several molecular properties were correlated with both the AP and ML axes of the bulb.

Given prior anatomical and physiological results^54-56^, we hypothesized that such correlations could arise, at least in part, due to similarity in the average response spectra of groups of sister cells receiving inputs from the same glomerulus. Because our individual FOVs are relatively small (~300-500 μm), we expect to encounter groups of co-glomerular sister cells that are similar in their average odor tuning. If, for example, two of such sister cell groups are on opposite sides of a field of view, we may observe an enrichment in the correlation between the cells’ locations and tuning, arising just from the difference in the average responses of these two sister cell groups.

To identify putative sister cells, we clustered cells on the basis of similarity in their responses to odors (Methods, Supplementary Figure 6, Supplementary Note 2). Clustering was carried out within and across FOVs, since sister M/T cells receiving primary input from the same glomeruli (as defined by their OR identity) are expected to be found on the dorsal aspect of the OB in multiple animals. The results of clustering are displayed using a t-sne projection^57^ in Figure 4c,d. After clustering, we collapsed the major clusters of cells into single ‘average’ cells, with both responses to odors and positions represented by the mean value within each cluster (Figure 4e,f). When this strategy was used for each FOV, all significant correlations between tuning to molecular properties and cells’ locations vanished. For example, for FOV#21 (Figure 4a,b,e,f), we plotted the correlations between the property response of the sampled tufted cells and their positions along AP and ML axes in a scatter plot (Figure 4g). Each point represents the spatial correlation of an individual molecular property. The properties significantly correlated (FDR q < 0.1) with either AP or ML bulb axes are displayed as blue and red dots. Before clustering (Figure 4g), we observed a large number of molecular properties correlated with either axis of the OB, while after clustering the putative co-glomerular T cells, we found none (Figure 4h).

We conclude that the correlations between M/T cells tuning to properties and their OB location are low and induced by the presence of groups of co-localized, similarly-tuned cells (presumably co-glomerular sister M/T cells). When such groups of M/T cells are collapsed together, the correlations between molecular properties and M/T cell locations over a spatial scale of ~0.5 mm disappear completely.

### M/T cells and glomeruli sample different molecular subspaces

We find that glomerular and M/T cell odor responses differ in their overall dimensionality, capture different amounts of variance with respect to tuning to the eDragon molecular properties, and display varying degrees of spatial correlation to these descriptors. Such discrepancies are consistent with a scenario in which the activity of M/T cells is shaped by extensive cross-talk between the OSN glomerular input and top-down cortical feedback and neuromodulatory action via local bulb circuits^37,40,58-62^. Therefore, we set to investigate further whether and how the odor spaces sampled by glomeruli and M/T cells differ from each other.

First, using the PCX method, we compared the percent variance in M/T cells and glomerular responses captured by the glomerular responses’ principal components (PCs) to the panel of 49 stimuli. If these two spaces were similar, the glomerular PCs would explain nearly the same fraction of variance in both the mitral cells and glomerular datasets. In contrast, glomerular PCs explained substantially less variance in the mitral versus glomerular responses (Figure 5a). Similarly, the PCs calculated for mitral cells odor responses were insufficient to describe the glomerular responses. However, the mitral cells PCs performed well in capturing tufted cell responses (Figure 5 a). Taken together, these findings suggest that the odor spaces probed by mitral cells and glomeruli are substantially different, while, within the granularity of our sampling, the tufted and mitral cells sample similar spaces.

**Figure 5.**
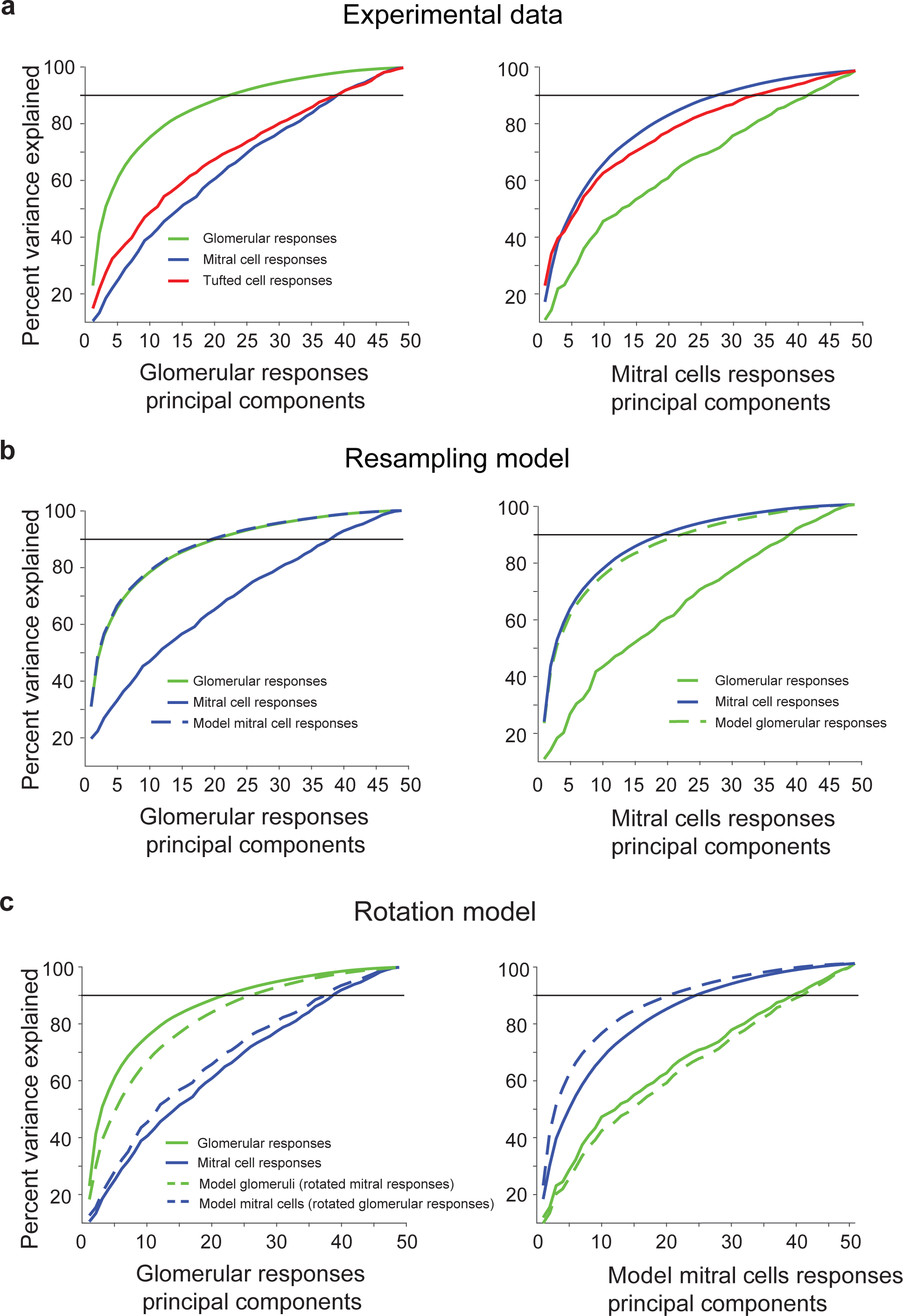
Comparison of glomerular and M/T cells odor response spaces. **a**. (*Left*) Percent variance explained of glomerular (green), mitral (blue) and tufted (red) cells responses shown as function of the number of included glomerular principal components, PCs. (*Right*) Percent variance explained of glomerular (green), mitral (blue) and tufted (red) cells odor responses shown as function of the number of included mitral cells responses principal components. **b**. Using a re-sampling model to explain the discrepancy between glomerular and mitral cell spaces. Model mitral cells responses were generated by taking samples from the glomerular responses. (*Left*) Percent variance explained of glomerular (green), mitral (blue) and simulated model mitral (dashed blue line) cells responses shown as function of the number of included glomerular principal components. Simulated mitral cell response variance (dashed blue line) appear similar to the explained glomerular variance (green) and deviate from experimentally observed mitral cell responses (solid blue). (*Right*) Percent variance explained of glomerular (green), mitral (blue) and simulated glomerular (dashed green line) responses shown as function of the number of included mitral cells response principal components. **c**. Model mitral cells responses were generated by rotating glomerular responses in odor space. Similarly, model glomerular responses were generated by rotating mitral cell responses. (*Left*) Percent variance explained of glomerular (green), mitral (blue), simulated model mitral (dashed blue line) cells and model glomerular (dashed green line) responses shown as function of the number of included glomerular principal components. (*Right*) Percent variance explained of glomerular (green), mitral (blue), simulated mitral cell (dashed blue line) and model glomerular (dashed green line) responses shown as function of the number of included model mitral cells response principal components.

One explanation for the differences in glomerular and mitral cell responses PC spaces is a sampling bias in probing the two layers, since our data includes a small population of mitral cells and glomeruli, albeit, from the same region of the bulb. In an attempt to reconcile the glomerular and mitral cell data, we tested a random selection model in which the responses of M/T cells reflect the responses of individual glomeruli. To obtain the responses of one M/T cell to the odors in our panel, we randomly selected a glomerulus in the dataset and assumed that neuronal responses faithfully relay inputs from this glomerulus. Within the random selection model, the discrepancy between glomerular and model mitral cell sampled spaces is small (green versus dashed blue lines, Figure 5b) and not compatible with the experimental data (green vs. blue, Figure 5a).

We sought to identify a model that could better explain the relation between glomerular and mitral cell odor spaces. We computed a rotation matrix in the odor space, which could convert the glomerular PCs to the mitral PCs space. More precisely, if 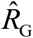 is the matrix of glomerular responses (glomeruli x odors), we generated surrogate mitral cell responses 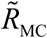 (mitral cells x odors) using the following equation:

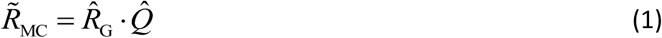

Here 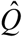is the rotation matrix 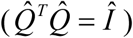that relates the glomerular and mitral cells PCs spaces. To obtain 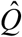, we used singular value decomposition, SVD. We call this transformation the rotation model, since it involves mixing glomerular responses to different odors with coefficients provided by rotation matrices to produce model mitral cell responses (Methods). Indeed, one can view the outcome of this procedure as rotating the glomerular PCs and projecting (squeezing and stretching) them on the mitral responses PCs. The mitral cells PCs may sample a subset of the glomerular PCs, as well as other (top-down) stimulus related information (i.e. expectation, behavioral value, etc.) which are underrepresented in the glomerular input. We find that the responses of model mitral cells (generated from glomerular responses, dashed blue lines, Figure 5c) are close to the variance produced by actual mitral cells responses (blue solid, Figure 5c), while those obtained from shuffled glomerular controls differ widely (Supplementary Figure 7). Thus, a simple rotation (Eq. 1) can generate mitral cell responses from glomerular responses and vice versa. Although these two dependencies are not overlapping completely, the discrepancies can be explained by noticing that glomerular responses, from which we generated the surrogate mitral cell responses, occupy a lower dimensionality PC space than the actual mitral cells (Figure 1g, h). For example, surrogate mitral cell responses in the model mitral PC space (Figure 5c, right) are left-shifted compared to the real cell responses because the dimensionality of glomerular PC space is lower (21D vs. 24D).

Our model described so far aims to relate the odor representations in the input and output layers of the olfactory bulb. The model described by equation (1) mixes glomerular responses to different odorants to yield M/T cell responses. Its goal is to relate glomerular and M/T response spaces using a minimum number of parameters. A circuit level model would be required to pool together inputs from multiple glomeruli (versus pooling single glomerular responses across odors) to produce M/T cell responses^63^. Such a model can be represented by a weight matrix multiplying the glomerular responses on the left rather than on the right as in equation (1):

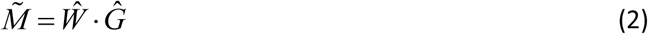

Here, 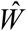 is the (glomeruli x glomeruli) weight matrix which is much larger than matrix 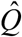. In the Methods section, we derive the weight matrix that yields exactly the same transformation as equation (1). Thus, equation (1) yields an equivalent, albeit more compact relationship between mitral and glomerular responses than equation (2).

Overall, we propose that the odor spaces of glomerular and mitral cell odor responses are non-trivially related via a rotation transformation. This transformation mixes responses obtained for different odors with predictable real-value coefficients, and may reflect the contribution of local processing and top-down centrifugal inputs impinging onto the mitral and tufted cells.

## Discussion

We imaged responses of populations of olfactory bulb inputs and outputs to a chemically diverse set of odors in awake naive head-fixed mice, and sought to relate this activity to the physical-chemical properties of odors. Our experiments show that odors activate glomeruli and M/T cells in a mosaic, spatially dispersed manner, with poor relation to an extensive set of physical-chemical molecular properties previously considered in the literature. Specifically, for both glomeruli and M/T cells, odors with similar physical-chemical descriptors did not elicit similar activity in the neuronal representations. The molecular properties were moderately predictive of the spatial location of glomeruli on a coarse spatial scale, but lacked predictive power for the M/T cells. Interestingly, even the large number of physical-chemical properties we considered was insufficient to explain the overall variance in neural odor responses of both the inputs and outputs of the olfactory bulb. Comparing activity patterns across the input and output layers, we found that glomeruli and mitral/tufted cells sample different stimulus subspaces, and identified a rotation transform in odor space that can relate these two sensory representations.

### Dimensionality of the olfactory bulb odor responses

There is wide disagreement on how many dimensions the odor space may have, ranging from several hundred, based on the number of odorant receptor types, to just 2-20 based on human perceptual judgments^7,22,23^. This is different, for example, from our understanding of color vision. There, three types of cones receptors form the 3 dimensions on which any neuronal and perceptual visual representations can be built^8^. In our experiments, the odor responses of a set of glomeruli and M/T cells from the dorsal aspect of the bulb to the 49 stimuli panel could be well-described within a ~20 dimensional principal components space (Figure 1). Systematically increasing the number of odors included in the analysis led to steady increase in the dimensionality of the observed responses. Thus, from these experiments, we can only extract a lower bound on the dimensionality of olfactory bulb neuronal odor representation. Future studies, probing much larger arrays of odors or optogenetic glomerular patterns in the bulb, as well as downstream olfactory brain areas, will help further constrain an estimate of the number of odor dimensions sampled by the rodent olfactory system.

We note that, overall, the responses of M/T cells are somewhat higher dimensional than those of glomeruli (24D vs. 21D). This could be a signature of M/T cells’ integration of lateral signals across glomeruli which are not available to optical imaging on the dorsal surface of the bulb, as well as of differences in sensitivity of the imaging methods used. In addition, local inhibitory inter-glomerular cross-talk, well as top-down cortical feedback and neuromodulatory input from multiple brain regions may add non-linearities and amplify the dimensionality of M/T cells responses. Such processing could enable comparisons across different OR input channels at the level of M/T cells, and help represent behaviorally relevant information in the bulb output.

### Mitral vs. Tufted cells

We examined the responses of M vs. T cells to the same set of odors at two different concentrations (~20 fold change). We found that T cells are more sensitive to odors, on average, than M cells, confirming previous observations^36-38^. We also found that T cells tend to generally increase activity for higher concentration of odors, but M cells could be modulated bi-directionally (Supplementary Figure 3, Supplementary Note 1). This is of interest because there have been suggestions that responses in higher olfactory areas such as the piriform cortex tend to be concentration invariant^64^, but OB responses tend to vary with concentration. Our data indicate that, at the population level, M cells, which send stronger inputs to piriform cortex than T cells^38^, could have much flatter response variation to concentrations, including some reduction in activity for higher concentrations. Such bi-directional, cell-specific dependence can arise from the known OB circuitry with extensive, distance-independent lateral inhibition^30,65,66^.

Despite these important differences in the responses properties of M and T cells to odors and concentrations, all conclusions about the representation of physical-chemical descriptors of odors and spatial layout are applicable to both populations.

### The relationship between molecular properties and neuronal responses in the bulb

We used the Principal Component Exchange (PCX) analysis to compare the PCA spaces of molecular properties and neuronal responses of both glomeruli and M/T cells to the same set of odorants. We found that the PCA spaces of properties and responses share very little overlap. This finding could be explained by the responses of both glomeruli and M/T cells including information that is not related to odor molecular properties *per se*. Such information may reflect valence, previous experience and expectations, or behavioral information, including changes in stimulus sampling^67^. Alternatively, it is possible that our set of 1,666 properties does not include additional relevant properties that also drive the responses of the olfactory neurons. Finally, neuronal responses may simply contain randomness that is unrelated to any useful signals, although the measured variability of responses to individual odors was less than 10%. Future experiments with mice engaged in different behavioral tasks, and additional chemical space characterization will help disambiguate these different possibilities.

### Can odor properties predict olfactory bulb odor responses?

Several studies have suggested that the molecular properties of odors can be used to predict the responses of neurons in the olfactory system. We addressed this question at the level of both OB inputs and outputs by asking whether similarity in molecular properties of pairs of chemicals can predict the similarity of neuronal populations to these same odors. Estimating the similarity of two chemicals was complicated by the fact that the large set of 1,666 molecular descriptors was highly redundant. Therefore, we used a sparsening procedure (non-negative LASSO) that imposes increasing costs with increasing number of used descriptors, such as to select those properties most predictive of the neuronal responses. Approximately 10 properties were sufficient to establish a substantial correlation (~0.60) between pairwise odor distances calculated in terms of molecular properties and neuronal response spaces for pairs of odorants. We tested the robustness of the relationship between molecular property similarity and neuronal response similarity in two ways (Figure 2, Supplementary Figure 4). First, for the M/T cells dataset, we iteratively selected sparse subsets of properties for one group of samples, and used these properties to predict responses for the remaining set of fields of view. This leave-out, cross-validation procedure yielded lower, but significant correlations between odor pairwise distances, implying that the properties predictive of responses generalize across samples – this is expected if similar regions of the OB are imaged across animals (for example in the glomerular layer). However, using a similar procedure, we found that properties predictive of similarity of neuronal responses to pairs of odors fail to generalize across new pairs of odors. Thus, the same sets of molecular properties could not be used to quantify odor identity distances in either glomeruli or mitral/tufted responses for different sets of odors and concentrations.

Our results differ from other published studies that have found predictable relations between molecular properties of odors and the responses of neurons in the early stages of the olfactory system ^14–16,19,68^. These differences could be due to several reasons. First, it is possible that due to the number of odors used (~50), the appropriate sets of properties were not established in a robust fashion. This seems unlikely since other studies have used similar number of odors, and did not systematically examine different concentrations of odors, as we did for the M/T cells. A second possibility, which we favor, is that there are correlations in the structure of neuronal responses, but these vary widely in terms of the molecular properties that are correlated. The LASSO procedure, because of the large number of fitting parameters, will always pick out some correlation between molecular descriptors and neuronal responses for a given training set. This fit, however, has very little predictive power for examples outside the training set. Previous work may suffer from this same problem of over fitting and poor generalization.

It is also important to note that relationship of olfactory neurons responses to these physical properties could be complex and highly nonlinear, and the algorithms used here may not capture it well, therefore encouraging the search for alternative analyses.

### Beyond a look-up table of physical-chemical properties (spatial order in the bulb)

There has been considerable debate on whether there is a continuous and recognizable map of chemical space in the olfactory bulb. Since microscopy offers spatial information inherently, we were able to ask whether the location of glomeruli or M/T cells has any relation to their selectivity to molecular descriptors of odorants. Significant correlation was observed in the glomerular layer (Figure 3) over a broad scale (~4-5 glomerular spacings). This is consistent with previous reports relating the odor spectrum and location of glomeruli^29,35,48,50^ which identified large chemotopic domains on the bulb surface (~1mm). The precision of these predictions is substantially lower (~6-8 fold) compared to the precision of the glomerular spatial layout across individuals (~0.5-1.0 glomerular spacings)^29^, but see^69^. Interestingly, several properties were significantly correlated along a preferred set of orthogonal axes (PC1 and PC2) rotated (~40 degrees, PC1) with respect to the AP-ML coordinates of the bulb. These may reflect specificity in the placement of odorant receptors (glomeruli) on the bulb surface during development, and are in agreement with the existence of different axon guidance gradients oriented differentially along the AP and ML axes of the bulb^27,31^.

While tiling the dorsal aspect of the bulb, the fields of view used for monitoring the M/T cells were smaller in comparison to the widefield imaging performed in the glomerular layer. This constrains our conclusions on spatial tuning of neurons to properties in the OB output layers to a finer scale (~0.5 mm). We observed weak, but statistically significant correlations between molecular properties and the location of somata in only a small number of fields of view. The correlations in such cases appear to be induced mainly by the presence of cells with highly correlated odor tuning, presumably because they are co-glomerular “sister” cells^54-56^. Since sister cells are spatially clustered^56^ (Supplementary Note 2), their presence within a field of view can increase the correlations between molecular properties and cell locations. For example, two large populations of putative sister neurons at the opposite sides of a field of view would create an appearance of spatial bias in the map. We removed such redundancy in responses, and replaced these groups of similarly tuned cells with single ‘average’ neurons represented by their mean odor response spectra and average positions within the group. This replacement resulted in the loss of any correlations between molecular properties and the spatial position of M/T cells (Figure 4). Thus, our results suggest that tuning to molecular properties included in the analysis, is moderately captured by the large spatial domains in the glomerular layer, but absent in the bulb output.

### Relating the glomerular and M/T cells odor response spaces

Although ideally this problem should be addressed by observing M/T and glomerular responses simultaneously in the same preparation, and with same activity sensors, obtaining such data was beyond the scope of the present study. Instead, we took advantage of data acquired in different animals to determine whether the spaces sampled by these two olfactory processing layers differ in a systematic fashion. We found that populations of glomeruli and M/T cells on the dorsal aspect of the bulb sample different subspaces with respect to the same panel of odors. In agreement, with previous work^37,40,58-62^, M/T cells do not simply relay the glomerular inputs to higher olfactory centers, but, in fact, substantially modify and diversify the OR input channels, as indicated by their higher dimensionality. Our data is consistent with a scenario in which glomeruli and mitral cell responses occupy intersecting, but distinct sensory odor spaces. Their different representations may reflect feedforward input from the olfactory epithelium, as well as local bulbar processing, and top-down cortical feedback and neuromodulatory input, which could sample information along different axes of the odor scenes. For example, mitral and tufted cells may filter out certain features of glomerular activity, but also integrate information that appears underrepresented at the level of the glomeruli. A rotation transform across the odor responses relates well these two spaces (Figure 5). This observation provides a potential framework for understanding the input-output function of the olfactory bulb in future studies aimed at probing simultaneously glomerular and M/T cell activity in naïve mice or during behavior.

### Relation to previous work

Our results contrast some previous reports that relate molecular properties of odors to neuronal responses in insects, fish, tadpoles and rodents^14-16,19,68^, but see^18,29,52^. These differences may arise due to several factors. Previous analyses^15,20,68^ have focused primarily on relating physical-chemical odor space to patterns of activity in the input layer (olfactory sensory neurons or glomeruli) in the anaesthetized preparations, while our study investigates population representations in the bulb inputs and output neurons in awake brain. Complex transformations in the olfactory bulb and top-down feedback^30,66^ could alter any relation between molecular properties and activity that present in the input layer. Differences could be accentuated in the awake brain – as discussed above, this could also contribute to the lower fraction of variance in neuronal data accounted for by the molecular features. There are also technical reasons for differences with previous reports. In particular, differences in the normalization strategies employed (for neural responses and for molecular properties), and a lack of cross-validation (both for odors and fields-of-view) may contribute to the overestimation of the predictive power of odor similarity metrics based on these molecular features for describing similarity of neuronal representations.

Importantly, we have focused here on the olfactory bulb activity and remain agnostic on the relationship between neuronal representations in downstream target brain regions and the physical-chemical properties. Future work, probing odor responses in cortical areas, will determine whether activity in higher olfactory centers is re-formatted such as to extract information on relevant molecular properties of odors.

Our data suggest that the physical-chemical properties used by us and others are not sufficient to fully represent the responses of OB neurons for the odors and range of concentrations sampled here. Indeed, these molecular properties poorly correlate with the glomeruli and M/T odor responses and account for only a small fraction of dimensions of the neuronal data (Figure 1). It seems that the molecular properties initially generated for computational chemistry studies do not capture stimulus features important for animals’ sensory perception, and novel descriptors are needed to link chemical space to neuronal representations. Relevant descriptors may carry information pertaining to behaviorally relevant properties of odors encountered by animals in their ecological niche as previously proposed (i.e. hedonic value, edibility, survival) ^7,11,22^ To date, the scarcity of reports on natural odor statistics for rodents has imposed constraints in the design of stimuli panels that faithfully sample the odor space encountered by rodents in their habitats. Systematic exploration of the natural odor statistics may offer further insight on the dimensionality and relevant features of odor space.

## Methods

### Chronic windows for awake head-fixed intrinsic and multiphoton imaging

Adult B6/129 and TBET-Cre X Ai38 GCaMP3.0 mice (males and females >80 days old, 25-40 g) were anesthetized with ketamine/xylazine (initial dose 70/7 mg/kg), supplemented every 45 minutes, and their heads fixed to a thin metal plate with acrylic glue. Heart beat, respiratory rate, and lack of pain reflexes were monitored throughout the procedure. Animals were administered dexamethasone (1 mg/Kg) to prevent swelling, enrofloxacin against bacterial infection (5 mg/Kg), and carprofen (5 mg/Kg) to reduce inflammation. To expose the dorsal surface of the OB for chronic imaging, a small craniotomy was made over both OB hemibulbs, using a either a biopsy punch ^70^ or thinning the skull with a 27 high-speed dental drill (Foredom, Bethel, CT), and removing it completely. A 3 mm glass cover slip (CS-3R, Warner Instruments) was placed atop and sealed in place using Vetbond (3M), further reinforced with cyanoacrylate (Krazy Glue) and dental acrylic (Lang Dental). A custom-built titanium head-bar was cemented on the skull near the lambda suture as described previously ^36–40^. Carprofen (5 mg/Kg) was administered for two days following surgery. Animals were left to recover for at least 48 hours after surgery before imaging and further habituated before the imaging sessions. All animal procedures conformed to NIH guidelines and are approved by Cold Spring Harbor Laboratory Animal Care and Use Committee.

### Odor stimulation

A custom odor delivery machine was built to deliver up to 165 stimuli automatically and in any desired sequence under computer control of solenoid valves (AL4124 24 VDC, Industrial Automation Components). Pure chemicals and mixtures were obtained from Sigma and International Flavors and Fragrances. Odorants were diluted 3,000 and respectively 100 fold into mineral oil and placed in blood collection tubes (Vacutainer, #366431) loaded on a custom made rack and sealed with a perforated rubber septum circumscribing two blunt end needles (Mcmaster, #75165A754). Fresh air was pumped into each tube via one needle by opening the corresponding solenoid valve. The mixed odor stream exited the tube through the other needle and was delivered at ~1l/min via Teflon-coated tubing to the animal’s snout. The concentration of the odors delivered to the mouse was measured using a photo-ionization device (PID; Aurora Scientific) and found to range between ~0.05-1 % saturated vapor pressure. The same PID was used to determine the time course of the odor waveform and the reliability of odor stimulation. A list of odors used in our experiments is provided in Supplementary Table 1. 49 out of the 57 stimuli used are monomolecular compounds, and were further included in the physical-chemical properties analysis. For the comparison of glomerular and M/T cell responses, the 1:100 dilution was used for same 49 odors. In a set of 6 M/T FOVs (3 mice), only first 33 odors in the panel were used.

In general, for intrinsic optical imaging (IOI) experiments, in each stimulus trial, we presented 12s of air, followed by 24 seconds of odor delivery. The interval between trials was at least 45s and each stimulus was repeated 4-5 times. Data was obtained from 6 mice (12 bulb hemispheres = hemibulbs). For two photon experiments, we scanned at 5-10 Hz/frame and covered a field of view up to ~350×500μm in the mitral cell layer. Before delivering odors, the OB was examined to gauge the quality of the surgery and select the regions of interest. The resting fluorescence of GCaMP3.0 is low, but could be discerned by frame averaging (~10 frames). Resting images at different depths were obtained before choosing specific optical sections for further experiments. Once a specific optical section was chosen, a time sequence of 120-240 frames was acquired. During the first 10 seconds, fresh air was delivered, followed by odor stimuli (of matched flow rate to the fresh air to avoid mechanical olfactory sensory neuron activation) for 4 seconds. Finally, fresh air was delivered for 10 seconds. The inter-trial interval was 45 seconds. Each odor was typically delivered 3-4 times. Mitral/tufted cell data presented here has been obtained from 19 distinct FOVs (15 mice). For 6 FOVs, the same panel of odors was presented at two nominal oil dilutions (1:3,000 and 1:100).

#### Intrinsic and multiphoton imaging

We used computer controlled LEDs to shine far red light (780nm) for imaging intrinsic optical signals on the dorsal surface of the bulb, acquiring images at 25 Hz (Vosskuhler, 1300-QF CCD camera). For two photon imaging, we used a Chameleon Ultra II Ti:Sapphire femtosecond pulsed laser (Coherent) and a custom-built multiphoton microscope. The shortest possible optical path was used to bring the laser onto a galvanometric mirrors scanning system (6215HB, Cambridge Technologies). The scanning system projected the incident laser beam tuned at 930nm through a scan lens and tube lens to backfill the aperture of an Olympus 20X, 1.0 NA objective. Scanning and acquisition were performed using custom Labview based software (National Instruments).

### Data analysis

All data matrices representing glomerular and M/T cells odor responses included in the analyses presented here are available upon request. For intrinsic imaging experiments, responsive glomeruli were identified as previously described^29^. For mitral and tufted cells, ROIs were manually selected based on anatomy. Care was taken to avoid selecting ROIs on cell bodies overlapping with neuropil (M/T lateral dendrites). To facilitate detection of responding cells or regions, we calculated a ratio image for each odor (average of images in odor period minus average of images in pre-stimulus period, normalized by the pre-stimulus average). We further obtained a maximum pixel projection of all odor responses, assigning to each pixel in the field of view the maximum response amplitude across the odor panel used - this allowed us to visually identify odor responsive regions. These responsive regions of interest mapped to individual mitral/tufted cell bodies respectively in the fluorescence image and were selected for further analysis ^29^

### Odor responses

To obtain odor responses, we computed the average florescence during the period of odor presentation for each trial *i, F*_2*i*_, and the average fluorescence during the preceding air period *F*_1*i*_. Responses were defined as statistically significant for p < 0.1 (2-way ANOVA). Because, in the subsequent analysis, additional more stringent statistical tests were performed, we reasoned that preliminary filtering of the data on the level of p<0.1 is reasonable. For the subsequent analysis, we used relative response (dF/F) defined as:

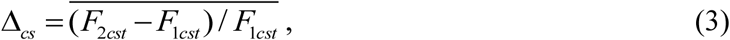

where indexes *c, s*, and *t* enumerate regions of interest (ROI, i.e. cells), odors, and trials respectively, while average is computed over the trials *i*.

For intrinsic optical imaging of glomeruli, we applied the same procedure as described previously^29^. Control ROIs drawn in non-responsive areas of the bulb were used to obtain a response signal threshold by comparison to odor responses in equal number of active glomeruli. Varying a signal threshold, the number of control ROIs that passed the threshold was compared to the number of responses in regions identified as active glomeruli to obtain a false positive ratio of <0.1. For both glomeruli and M/T cells, the ROI-odor pairs with non-significant response were set to 0.

### Hierarchical clustering

To identify mitral cell bodies with similar odor tuning, we performed a cluster analysis based on the similarity between odor response spectra (average linkage or complete linkage clustering, cutoff at similarity = 0.7). Each cluster with three or more members was taken to represent potential sister M/T cells receiving common primary input from the same parent glomerulus and considered for further analysis.

### Odor physical-chemical property response (PR) and property response spectra (PRS)

To determine the sensitivity of individual ROIs to various properties we evaluated the property response (PR) and property response spectra (PRS), the ROIs’ chemical receptive field, Φ_*cp*_. Each element of the matrix Φ_*cp*_ with indexes *c* and *p* indicates the strength of response of a given ROI, *c* to an odor physical-chemical property, *p*. Thus matrix Φ_*cp*_ describes the physical-chemical tuning of the set of ROIs. To compute Φ_*cp*_, we first calculated the values of 1,666 physical-chemical properties (PCPs) for the odors used in out panel. This resulted in a property matrix *P*_*sp*_. *S* enumerates the monomolecular odors (1-53 for M/T cells and 1-49 for glomeruli), as above, while the second index *p* ranging between 1 and 1,666 denotes the PCPs. The property matrix, was obtained by downloading molecular structures for the monomolecular odors from PubChem and using an online set of algorithms eDragon to evaluate the PCPs. The full list of molecular descriptors (PCPs) used can be found here: http://www.talete.mi.it/products/dragon_molecular_descriptor_list.pdf.

Because the properties obtained were highly inhomogeneous in ranges and scales, we normalized the data as outlined below. If a property took both negative and positive values for the odors in the panel, we subtracted the mean value for this property and divided by the standard deviation across the odors in the panel. If a property was strictly positive (molecular weight, etc.), we examined the standard deviation of its logarithm (SDL). If SDL was larger than 1, we assumed the property is lognormally distributed. The corresponding property was replaced in matrix *P*_*sp*_ by its logarithm with the mean subtracted. If SDL was smaller than 1, we subtracted the mean from the property and divided the values by its standard deviation for the odors in the panel. Normalizing each property by the standard deviation, we minimized the influence of measurement units on the dynamic range of some properties which could introduce a subjective bias. By implementing this procedure, all properties were brought to zero mean and similar standard deviations. The resulting matrix was denoted 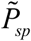. ~200 physical-chemical properties did not take non-zero values for the odors in our panel and were not included in the analysis.

To compute the property response spectrum (PRS), the ROI’s chemical receptive field across the properties, we used the following formula:

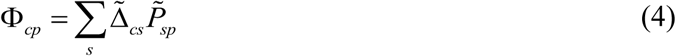

For each region of interest (ROI) *c*, a property response (PR), defined as an entry in the PRS, was equal to the correlation between the dF/F odor responses of this ROI and the property strength vector (PSV), given by the values taken by property *p* computed over the entire set of odors presented (enumerated by *s*).

To evaluate the significance of the correlation between a property and a neuron’s response pattern, we use a p-value threshold of.05. p-values are calculated using MATLAB’s built-in function corr(). This implementation evaluates the t statistic as 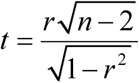, where *r* is the correlation coefficient and *n* is the number of data points. The p-value is then twice the probability a t distributed variable exceeds *t*.

### Odorant Response Similarity and Property Space Similarity

We compute the correlation matrix of the relative responses (dF/F) ∆_*cs*_ and ∆_*ck*_ between each pair of odors, s and k, for all glomeruli or cells, c as:

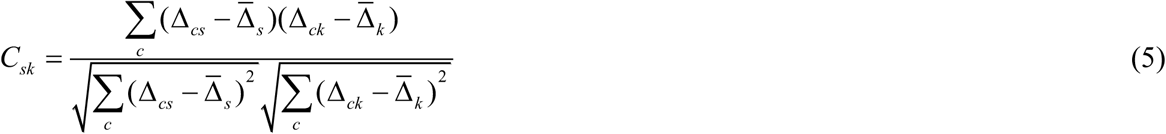

Where 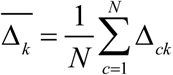. Indexing the unique pairs of odors of *C*_*sk*_ with i, we have a vector of correlations, 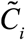. Using the property matrix 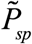 described above, we calculate the Euclidean distance matrix as:

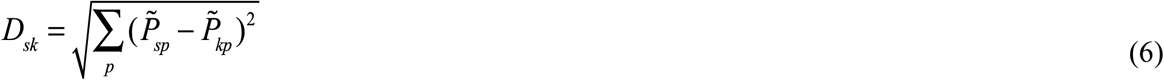

We calculate Euclidean distances in neuronal response space between each smell s and k as 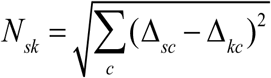. 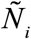 is the vectorized unique smell pairs of *N*_*sk*_. Figure 2a,b (*Top*) plots 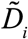 against 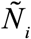. Following the same vectorization procedure as with the response correlations 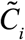, we vectorize the unique pairs of odors in matrix *D*_*sk*_ with 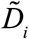. Figure 2a,b (*Bottom*) shows a plot of 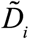 versus 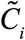.

We searched for a subset of properties which generate property distances 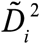 that correlate best with neuronal response distances 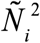. For this, we used the LASSO algorithm ^45^. This algorithm minimizes the following equation :

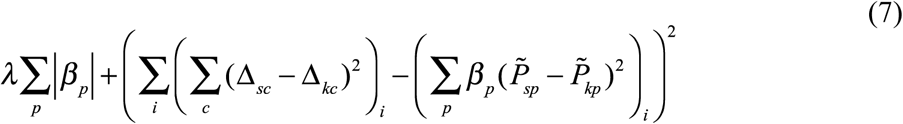

Where ( )_*i*_ denotes the vectorization of the unique pairs of odors s and k described above. Manipulating each weight parameter *β*_*p*_ (non-negative) to minimize the term 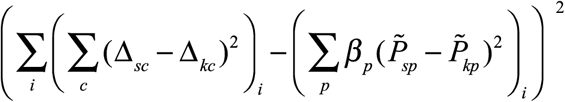 creates a property space with square property distances 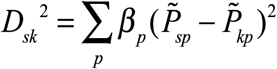 that reconstruct the square neuronal response distances 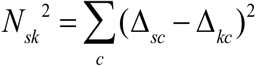. The first term of the LASSO objective function, 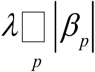, penalizes the use of nonzero weights (Supplementary Figure 4f). This forces the algorithm to choose the most parsimonious property space to reconstruct neuronal response space. Increasing the parameter *λ* puts more pressure on each *β*_*p*_ to be zero. Figure 3c,d shows, for different number of non-zero properties, the correlations between distances in the weighted property space 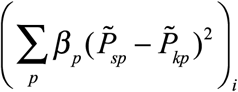 and distances in the neuronal response space 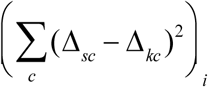.

To cross-validate on new responses, we randomly selected and withheld half of the FOVs and performed LASSO regression on the remaining data. Then, we recomputed the correlations between property distance and response distance in the withheld data.

To determine how our results generalized for new stimuli, we removed one pair of odors and performed the above analysis on the rest of the data. Then we calculated the distance between the two removed odors in the reduced property space found by LASSO regression. We repeated this procedure for each pair of odors i, with each repetition generating one property distance 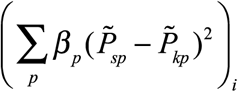 Then, we found the correlation between the vector of these property distances and the neuronal property distances 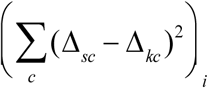.

Finally, we cross-validated with new FOVs and new odors by combining the two previous crossvalidation procedures. That is, we withheld one pair of odors and half of the FOVs, and then performed LASSO. Using the reduced property space found by LASSO, we then predicted the response distance of the two removed odors in the withheld response data.

The physical-chemical properties selected by LASSO regression were different for crossvalidated and non-cross-validated curves in Figure 2. This is because data available in each case is different. For FOV cross-validation, a different set of properties emerged for each subset of selected FOVs in the training set. Similarly, for odor cross-validation, the distance between each pair of odors was calculated using a different data subset and led to distinct sets of best properties.

### Greedy algorithm

First, we found the property for which the Euclidean distance of each pair of odorants is best correlated with the odor response similarity, defined as the correlation between the M/T neuronal response profiles (cell response spectra, CRS) in all FOVs of these two odorants. Second, we searched through all remaining properties and with each iteration added to the metric the property which most greatly increased the correlation between the physical-chemical property distance and neuronal response odor similarity. The process terminated once adding any new property decreased the correlation (Supplementary Figure 4g).

### Correlations between ROI property response spectra and ROI locations

To evaluate PCP-to-position correlations, we computed the Pearson correlation coefficients between the locations of ROIs 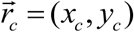 and their property response spectra (receptive fields, Φ_*cp*_). Here *x*_*c*_; and *y*_*c*_ are AP and ML positions of an ROI *k* on the surface of the bulb. If the average locations are 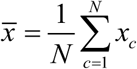 and 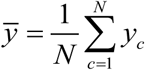, the Pearson correlation for a property *p* with the position of the ROI along the AP axis is defined as:

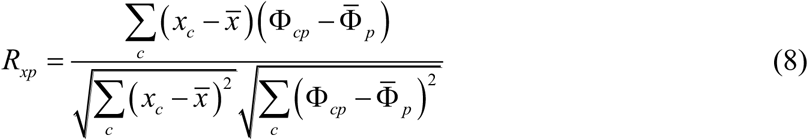

Where 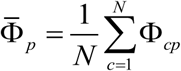. Similarly the correlation of the PCP-to-ML position, denoted as ‘y’ correlation is defined as:

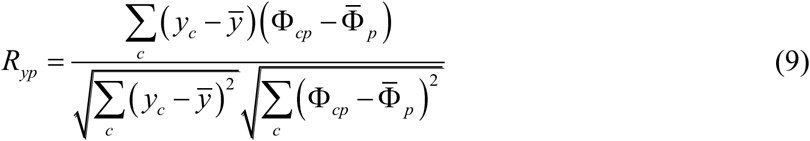

In addition to calculating the correlation values, we evaluated their statistical significance using MATLAB function ‘corr’. The corresponding p-values were computed for each property: *P*_*xp*_ and *P*_*yp*_ We applied False Discovery Rate correction^53^ to each set of p-values and found the set of q-values *Q*_*xp*_ and *Q*_*yp*_ using the MATLAB function ‘mafdr’. A property was assumed to be significantly correlated with A-P or M-L axes if the corresponding q-value was less than 0.1. The properties with significant correlations are shown in Figure 4a,b by color.

### Predictions of glomerular position based on odor physical-chemical properties

We tested whether cells’ tuning properties are predictive of their locations in the bulb. To this end, for each field of view and for each bulb axis, we built a linear regression that should approximate the positions of glomeruli, c:

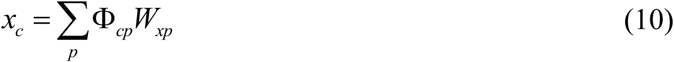

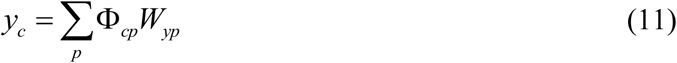

where *W*_*xp*_ and *W*_*yp*_ are sparse vectors of unknown coefficients that were found using the LASSO algorithm^45^. We added a column of ones to the matrix Φ_*cp*_ to include a possible offset to the approximation of coordinates. We ensured vectors *W*_*xp*_ and *W*_*yp*_ have only 20 non-zero components. To validate the prediction built on the basis of glomerular receptive fields, we excluded a single ROI (glomerulus) from the dataset, further obtained regressions with the LASSO algorithm based on the remaining ROIs, and then used the removed glomerulus to test the quality of prediction. We repeated this procedure for all glomeruli in the dataset. The resulting predictions for glomerular positions were compared to the actually observed bulbar positions and the quality of predictions was evaluated by computing the distance between actual and predicted positions measured in terms of average glomerular size (AGS, Figure 3g,h).

#### PCA space comparison using PC exchange (PCX) method

To compare the odor spaces sampled by mitral cells versus glomeruli, we projected them onto each other’s principal components. More explicitly, we consider the matrix of mitral cell responses, with element *M*_*ms*_ corresponding to mitral cell *m* responding to odor *s*, and similarly the response matrix with elements *G*_*gs*_ for glomerulus *g* and odor *s*. For each response matrix, singular value decompositions can be written as:

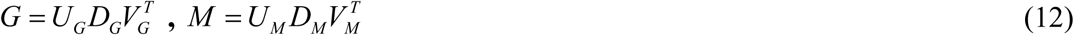

where *U*_*G/M*_, *D*_*G/M*_, and *V*_*G/M*_ are the unitary matrix, diagonal singular value matrix, and eigenvector (PC) matrix respectively for the glomerular and MTC response matrices. If dimensions of the matrix *G* are [*N*_*G*_ × *N*_*s*_], i.e. number of glomeruli by the number of odors, the dimensions of matrices *U*_*G*_, *D*_*G*_, and *V*_*G*_ are *N*_*G*_ × *N*_*s*_, *N*_*s*_ × *N*_*s*_, and *N*_*s*_ × *N*_*s*_ respectively. Here the number of odors is smaller than the number of glomeruli. We then computed projections of mitral cell responses onto the glomerular principal components, *M*_*G*_, and the projections of glomerular responses on the mitral cell principal components, *G*_*M*_, as follows:

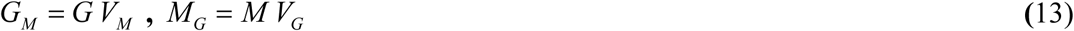

The variance of the glomeruli projections for each mitral cell principal component was calculated as:

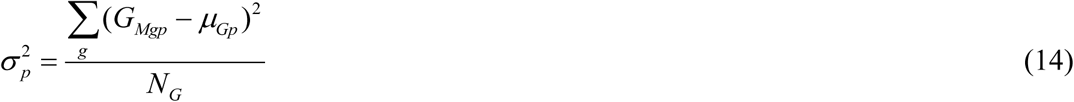

Where *μ*_*Gp*_ is mean of the glomerular responses on the mitral cell principal component *p* and *N*_*G*_ is the total number of glomeruli. Variance for mitral cells projected onto glomeruli can be found by replacing every instance of G with M and g with m.

The random data control used in Figure 1 is a random matrix, *N*_*G*_ × *N*_*s*_, constructed by sampling from a Gaussian distribution centered at 0 with SD = 1.

#### Resampling model

In this model, we tested whether mitral cell responses are a re-sampling of glomeruli responses. We randomly selected glomeruli with repetition to generate a new response matrix with equal number of cells as the true mitral cell response matrix. We then used the PCA space comparison method described above to compare the resampled glomeruli to the true mitral cell responses.

#### Rotation model

we tested whether mitral cell responses constitute a rotation of the glomeruli’s sampling of odor space. Using the notation from the PCA space comparison method, we modeled surrogate mitral cell responses to be:

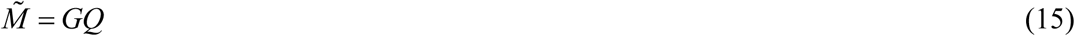

Where *Q* is a *N*_*s*_ × *N*_*s*_ rotation matrix, calculated as:

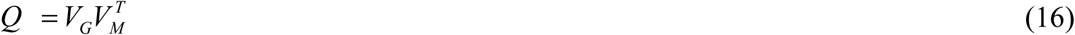

Because 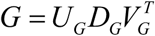 and 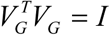, for the surrogate mitral cell responses (15) we obtain 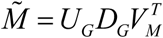. We further compared the rotated glomeruli response matrix with the mitral cell response matrix using the PCA space comparison method described above.

Rotation matrix *Q* cannot be viewed as a connectivity matrix. This rotation does not make predictions regarding the specific connectivity of individual glomeruli and mitral cells. Instead, it enables us to compare the two layers’ sampling of odor space. This is because, in equation (15), matrix *Q* multiplies the glomerular responses *G* on the right thus mixing responses of glomeruli to different smells to obtain the surrogate mitral cell responses, 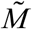. To obtain a weight matrix *W* that mixes glomerular responses for the same odor to obtain the same surrogate matrix, 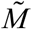 one would have to multiply the glomerular matrix on the left, i.e.

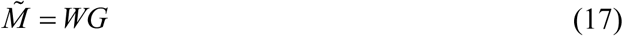

Because 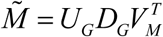 and 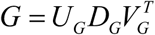, the weight matrix can be identified:

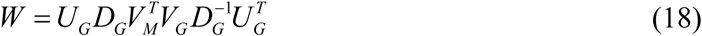

These two equations can be viewed as feedforward network equations that produce mitral cell responses from glomerular activities.

